# A pan-cancer gamma delta T cell repertoire

**DOI:** 10.1101/2024.07.18.604205

**Authors:** Xiaoqing Yu, Li Song, Ling Cen, Biwei Cao, Ranran Tao, Yuanyuan Shen, Daniel Abate- Daga, Paulo C. Rodriguez, Jose R. Conejo-Garcia, Xuefeng Wang

**Affiliations:** Department of Biostatistics and Bioinformatics, H. Lee Moffitt Cancer Center & Research Institute, Tampa, FL 33612, USA; Moffitt Cancer Center Immuno-Oncology Program, Tampa, FL 33612, USA; Department of Data Science, Dana-Farber Cancer Institute, Boston, MA 02215, USA; Department of Immunology, H. Lee Moffitt Cancer Center & Research Institute, Tampa, FL 33612, USA; Department of Integrative Immunology, Duke University, Durham, NC 27710, USA; Current: Department of Biomedical Data Science, Geisel School of Medicine, Dartmouth College, Lebanon, NH 03756, USA

## Abstract

This report presents the largest collection of gamma-delta T cell receptor (γδ TCR) reads in human cancer to date, analyzing about 11,000 patient tumor samples across 33 cancer types using the TRUST4 algorithm. Despite γδ T cells being a small fraction of the T cell population, they play a key role in both innate and adaptive immunity. Our comprehensive analysis reveals their significant presence across all cancer types, specifically highlighting the diverse spectrum and clonality patterns of their γδ receptors. This research highlights the complex roles of γδ T cells in tumor tissues and their potential as prognostic biomarkers. We also demonstrate the utility of T cell receptor gamma (TRG) and delta (TRD) gene expression values from standard RNA-seq data. Ultimately, our work establishes a fundamental resource for future tumor-infiltrating γδ T cell research and may facilitate the development of novel γδ-T-cell-based therapeutic strategies. Together, we demonstrate the strong diversity and prognostic potential of γδ T cells in multiple cancer types.

**Highlights:** Comprehensive analysis of γδ TCRs from 11,473 tumor samples

Significant variability and overall consistency in γδ gene expression and clonotype

γδ TCR expression and diversity as prognostic biomarkers across multiple cancers

Centralized γδ TCR repertoire database for future therapeutic discovery

## INTRODUCTION

Gamma delta T cells (γδ T cells) represent a specialized subset within the broader T cell population, distinguished by their T cell receptors (TCRs) that pair one γ-chain with one δ-chain. Like other T lymphocyte, the diversity of γδ T cells’ TCRs is realized through V(D)J recombination, a process that randomly rearranges variable (V), diversity (D), and joining (J) gene segments. While constituting only a small portion of the overall T cell population, they are an important subset that uniquely contributes to both innate and adaptive immunity. Unlike their alpha beta (αβ) T cell counterparts, the mechanism underlying antigen recognition by γδ T cells is not yet fully elucidated^1^. The γδ TCRs have significantly fewer number of variable (V) genes compared to α and β chains. Yet, it is evident that they can recognize antigens in a major histocompatibility complex (MHC)-unrestricted manner, enabling them to respond to a broad range of pathogens and malignant cells^2^. During recent years, there has been an explosive research interest in γδ T cells in the realms of cancer immunology and immunotherapy, owing to their unique regulatory roles and potential antitumor functions. γδ T cells, especially the Vδ1 subpopulation, have been found to infiltrate most tumors^3^. On the other hand, circulating γδ T cells in peripheral blood and lymphoid tissues are composed of different subpopulations, with a predominant subset expressing the Vδ2 receptor^4^. Specific subpopulations, such as Vγ9Vδ2, play a critical role in immune surveillance against infections and tumorigenesis, and are present in both peripheral blood and tumor tissues. γδ T cells are also known for producing various cytokines and chemokines, such as interferon gamma (IFN-γ) and tumor necrosis factor alpha (TNF-α). In addition, they are instrumental in a range of activities: from regulating B cells and dendritic cell maturation to recruiting macrophages and exhibiting cytolytic activity against various cancer cells^5–8^. Recent research on both human and murine samples has revealed functional plasticity in γδ T cells: the majority of γδ T cells in pre-malignant or non-tumor tissues demonstrate cytotoxic effects, whereas certain tumor-infiltrating δ T cell subsets can display pro-tumorigenic effects^9^. A more recent breakthrough discovery indicates that the antitumor efficacy of γδ T cells can be potentiated or restored through targeted therapeutic approaches, specifically by targeting BTN3A1 to activate Vδ2 T cells^10^. More recently, γδ T cells were found associated with response to immune checkpoint inhibition in cancers with low HLA expression^11^. These discoveries open a potential new avenue for treating cancers that are unresponsive to existing immunotherapy interventions.

Despite the recognized importance of γδ T cells in both functional and translational research, a comprehensive understanding of their repertoire within tumor tissues remains elusive. New single-cell technologies like the 10X Chromium single-cell V(D)J enrichment solution have shed light on the diversity and clonality of T cells within tumors. However, these technologies primarily target the αβ chain TCR and face limitations in sequencing depth, cost, and scalability of samples. In contrast, if repurposed for immune receptor repertoire analysis, bulk tumor sequencing offers a holistic approach for the tumor microenvironment assessment, capturing a wide spectrum of γδ TCRs. Many computational tools^12–15^ have been developed to perform de novo assembly of the complementarity determining region 3 (CDR3) sequences using blood and tissue RNA-seq data. Among them, TRUST^13,14^ stands out as a sensitive and reliable method for detecting TCR CDR3 sequences from tumor RNA-seq data. Its distinct approach, which involves pairwise read comparisons, enhances the detection of the less abundant TCR clones. The results from these tools have become instrumental in elucidating the complexities of tumor-infiltrating T cell repertoires. For instance, by analyzing The Cancer Genome Atlas (TCGA) tumor RNA-seq data, TRUST was able to identify the full spectrum of T cell and B cell repertoires that infiltrate tumors across cancer types, shedding light on their probable roles in tumor immunity^16^. Recently, more enhanced features have been incorporated into the upgraded algorithm, called TRUST4, which has shown better benchmarked performance^17,18^. These computational advancements offer opportunities for addressing the significant knowledge gap concerning the less studied γδ T cell repertoire—via revisiting pan-cancer datasets.

In this report, we reanalyzed raw RNA-seq data from nearly 11,000 tumor samples utilizing the latest version of TRUST4. This allowed us to offer the most accurate and comprehensive examination of the γδ TCR landscape spanning 33 different cancer types, making this the largest collection of γδ TCR reads in human cancer to date. A series of downstream analyses were performed to further investigate the pan-cancer repertoire landscape, the association between γδ TCR clones and VDJ gene expression, as well as their implications in patient prognosis. Collectively, our results and findings serve as a foundational resource for γδ T cell research in oncology.

## RESULTS

### Reconstructing gamma-delta TCRs (**γδ**-TCRs) from 11,473 tumor RNA-seq samples

We utilized the computational method TRUST4 to process RNA-seq data, encompassing 10,970 BAM files, from 10,131 patients within the TCGA database (Figure 1). The vast majority of these samples consist of paired-end RNAseq data, except for 637 samples which are single-end. This updated immune repertoire reconstruction tool allowed for the efficient and sensitive de novo assembly of CDR3 sequences derived from TCR transcripts. Through this computationally intensive effort, our objective was to curate the most comprehensive pan-cancer catalog of γδ-TCRs across all cancer types represented in TCGA. All analyses presented in this study were based on the TCGA TCR assembly data that was fully completed in January 2024. From the entire TCGA RNA-seq dataset, we extracted approximately 3 million reads covering the TCR CDR3 region. These reads were annotated with information such as amino acid sequences and associated variable (V) and joining (J) genes. To reduce false positive calls, we applied a series of stringent filtering criteria—by removing biologically implausible and ambiguous sequences, e.g., unproductive CDR3s and those concurrently annotated as αβ-TCRs. This process led to the identification of 34,129 unique γδ-TCR clones. Among the analyzed tumors, 6,751 tumors yielded at least one copy of a γδ-TCR clone. In total, when considering only one tumor sample per patient, we identified 31,924 unique clones. After further filtering to exclude out-of-frame CDR3s, this number was refined to 22,205 unique clones (Figure 1).

**Figure 1.**
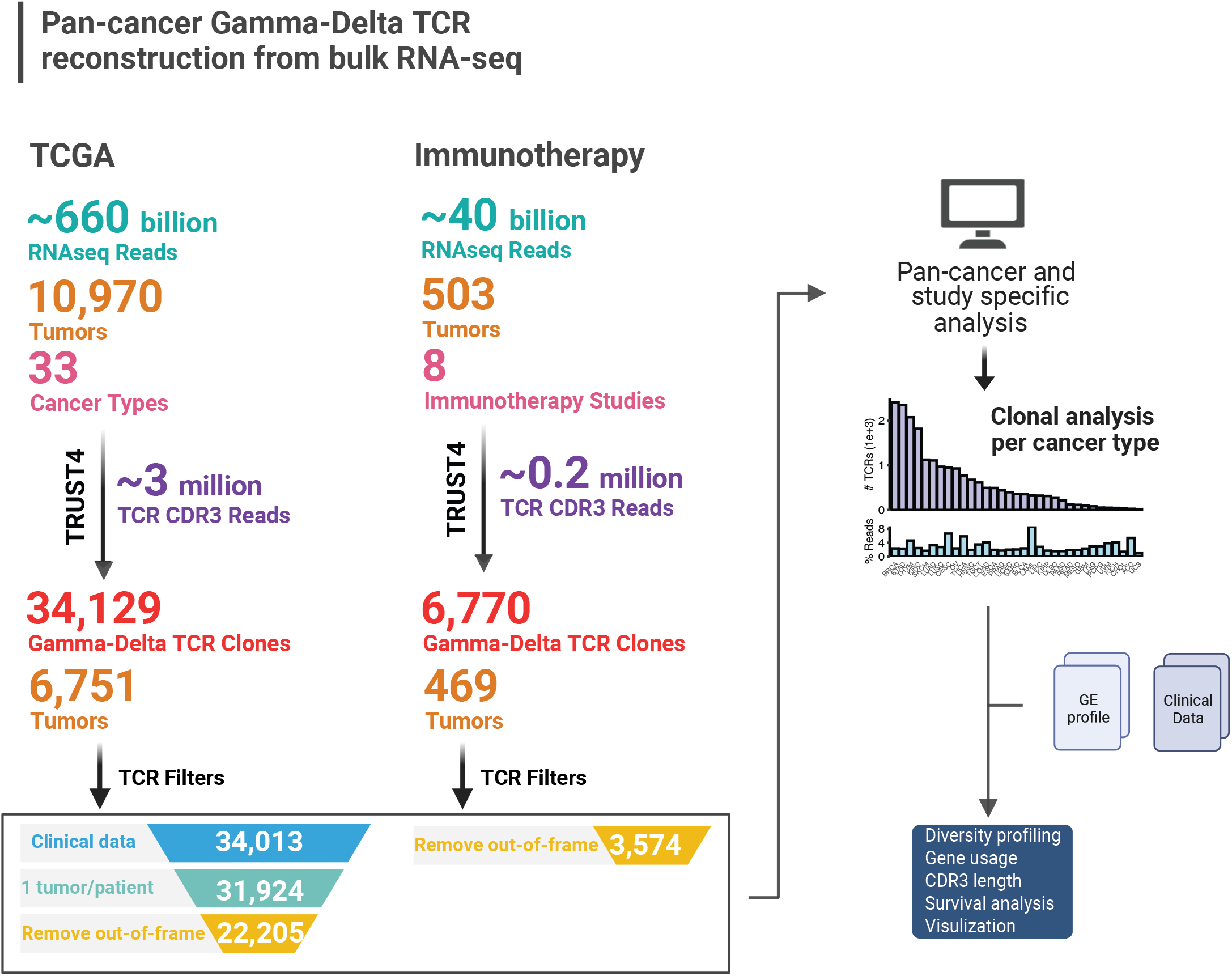
Comprehensive pan-cancer analysis of gamma-delta TCRs. 660 million RNA-seq reads from a total of 10,970 tumor samples in the TCGA dataset were reanalyzed for gamma-delta TCRs using TRUST4. It led to the identification of 34,129 unique γδ-TCR clones of 6,751 tumors. γδ-TCRs were further filtered to remove 1) samples without clinical information, 2) metastasis tumors for patients with multiple tumors available, 3) out-of-frame CDR3 sequences as well as CDR3 sequences carrying alpha-beta V genes. Similarity, approximately 40 million RNA-seq reads from 503 tumors downloaded from 8 immunotherapy studies were reanalyzed, generating 6,770 unique γδ-TCR clones, of which 3,574 clones from 459 tumors were retained for downstream analysis. Comprehensive pan-cancer and cancer/study specific γδ-TCR repertoire analysis was conducted using the filtered γδ-TCR sequences. These refined data sets, representing a significant resource in γδ-TCR research, were made available to the research community, facilitating broader scientific exploration.

In addition to the TCGA pan-cancer dataset, we also downloaded and processed raw RNA-seq data of a total of 503 tumors from 8 immunotherapy studies (see Methods). Similar to the TCGA dataset, we performed TRUST4 analysis and extracted approximately 0.2 million reads covering the TCR CDR3 region. This process led to the identification of 6,770 unique γδ-TCR clones.

### A pan-cancer catalog of **γδ**-TCRs reveals significant variability across cancers

We explored the Gamma chain expression and clonotype count information across 33 distinct cancer types (Figure 2A). TCR gamma gene expression for each sample is represented by an enrichment score, calculated based on ssGSEA on all TCR gamma genes (e.g., TRGV9). We observed significant variability in enrichment scores, both between and within cancer types (Figure 2A top panel). As expected, tumors traditionally classified as immune-cold cancers^19^, such as UVM and GBM, demonstrate the lowest enrichment scores, while lymphatic system-related cancers like LAML, THYM, and DLBC have the highest. Beyond THYM, the top five solid-tumor cancer types carrying the highest enrichment are PRAD, KIRC, TGCT, LUAD, and KICH. We next examined the number of patients that have at least one copy of a TCR-gamma clone and the number of patients with no reconstructed functional TCR-gamma clone for each cancer type (Figure 2A second panel). In cancers that rank highest in gamma gene enrichment (spanning from LAML to STAD), cancers generally have a high proportion of patients with detectable TCR-gamma clones, with PRAD being a notable exception.

**Figure 2.**
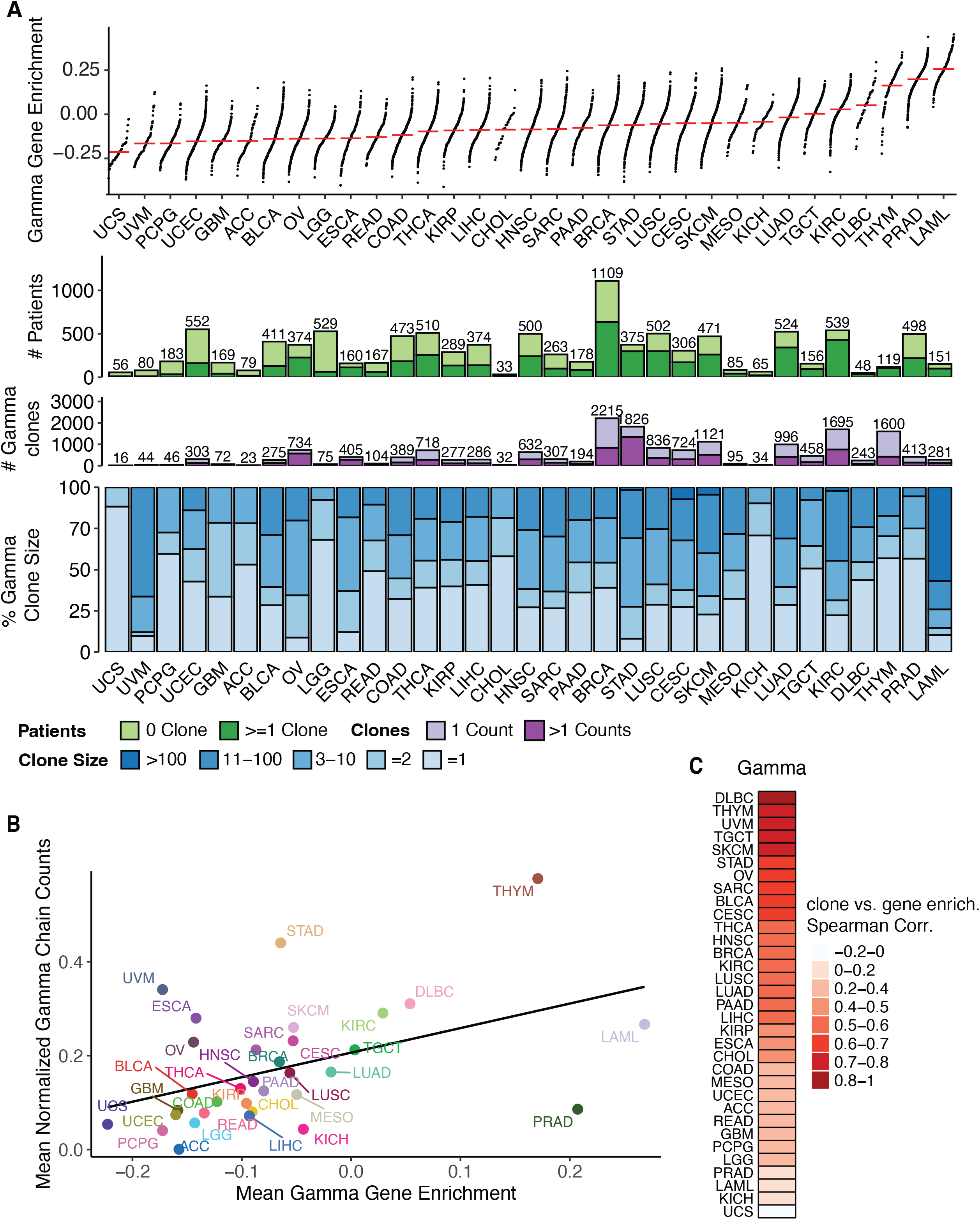
TCR gamma chain expression and clonotype distribution across TCGA cancer types. (A) Variability in TCR gamma gene enrichment and clonotype presence across 33 distinct TCGA cancer types. Top panel shows gamma gene enrichment scores of tumors. Each dot represents a tumor, with y-axis position indicating the normalized gamma gene enrichment score. Cancer types are ordered by their median enrichment scores, with the lowest scores on left and highest on right. The second panel shows the number of patients in each cancer types, with light green indicating patients without any γ TCR identified and dark green indicating patients with at least one γ TCR clone identified. The third panel shows number of unique γ TCR clones identified in each cancer type, with light purple representing clones with only 1 read and dark purple representing clones with > 1 reads. The bottom panel shows the proportion of unique γ TCR clones with specific clone sizes: =1 read, =2 reads, 3-10 reads, 11-100 reads, and >100 reads. (B) Correlation between TCR gamma gene enrichment scores and normalized gamma chain counts across cancer types. Each dot represents a cancer type with y-axis indicating the mean normalized gamma chain counts and x-axis indicating mean gamma gene enrichment scores across tumors. (C) Spearman correlation between gamma gene enrichment scores and normalized gamma chain counts within each cancer type, with correlation coefficients shown in different gradients of red.

Among the intermediate-ranked cancers, such as BRCA, HNSC, and THCA, approximately 50% of patients have detectable gamma clones. The lower-ranked cancers follow expected trends of fewer clones, but with exceptions like OV. We investigated the number of TCR-gamma clones along with the clone size distribution across cancer types (Figure 2A third and bottom panels). Overall, a dominant presence of large clone segment is not observed in most cancers. Cancers including LAML, KIRC, LUAD, SKCM, CESC, LUSC, STAD, SARC, HNSC, ESCA, OV, and BLCA do display a relatively higher proportion of intermediate to larger clones compared to other cancer types. An analysis of delta gene expression and clones reveals a similar pattern (Supplementary Figure S1). Compared to their gamma gene counterparts, delta gene expression and clone frequency are notably sparser across various cancer types. It is important to note that these clone sizes should be interpreted as down-sampled relative frequencies, considering the small proportion of γδ-TCRs and the fact that we are repurposing bulk RNA-seq for immune repertoire analysis. The actual clone sizes are anticipated to be much larger when profiled using a targeted gamma chain TCR assay.

We further checked the distribution of out-of-frame TCR reads (Supplementary Figure S2).

Although out-of-frame (OOF) TCRs do not contribute to functional immune response, the ratio of in-frame/out-of-frame TCRs could serve as a novel biomarker, and these sequences might provide insights into the diversity of progenitor T cells. However, we did not observe any cancer-specific patterns in terms of in-frame/out-of-frame ratios (Supplementary Figure S2A). The range of these ratios in gamma chain was rather uniform across different cancer types, varying between 27% to 41%. The overall distribution of OOF clone abundance mirrored that of in-frame reads, with the highest yields observed in STAD and BRCA. Most of these clones are singleton counts, suggesting that they are indeed non-functional. A similar pattern was observed in TCR delta chains, albeit with a significantly sparser distribution.

In our data, the CDR3 length distribution of γδ TCRs is in line with the patterns in beta chains^13^, where the lengths of these CDR3 regions predominantly range from 10 to 15 amino acids (Supplementary Figure S3). Overall, the delta chain tends to have a slightly longer region compared to the gamma chain, which could be due to the inclusion of D genes. When we rank cancer types by the length of the CDR3, there is a noticeable inconsistency between the rankings for gamma and delta chains.

### Overall consistency between **γδ** gene expression and clonotype

Next, we conducted a comparison between the mean Gamma gene enrichment score with the mean normalized gamma chain counts across the 33 cancer types, and observed an intermediate positive correlation pattern (Figure 2B). Several cancers, such as STAD and THYM, are observed to slightly deviate above the trend line, suggesting higher TCR-gamma reads than what might be expected from the gene expression-based enrichment score. Cancers such as BRCA, TGCT, HNSC, and BLCA align closely to the trend line, indicating a more direct relationship between their gene expression and TCR-gamma readouts. Conversely, PRAD stands out as a notable outliner below the trend line, emphasizing that the gene expression might not always mirror the total abundance or diversity of productive CDR3 sequences. This pattern highlights the complexity of the relationship between gene transcription levels and the resulting functional immune repertoire. For a more in-depth sample-wise analysis, we further examined the relationship between normalized gamma CDR3 counts and both gamma and delta gene expression enrichment within each cancer type, accounting for tumor purity (Supplementary Figure S1B). About two-thirds of the cancers exhibited intermediate to strong correlations, confirming a generally predictable relationship between gene expression and TCR-gamma reads for the majority of cancer types examined. In addition to the normalization chain counts, we calculated the clonality metrics for all cancer types (Supplementary Figure S4). Our analysis revealed a broad spectrum of clonality, with values ranging from 0 to 1 (larger values mean more clonal expansion), across patients in all cancer types. Furthermore, it was noted that the most immunogenic cancers, specifically SKCM, BLCA, and HNSC, generally display higher median clonality.

### **γδ**-TCR V-J gene pair distribution across cancer types

The landscape of γδ-TCR V-J gene pair distribution offers another layer of insight into clonality and the dynamics of γδ immune repertoire across different cancer types (Supplementary Figure S5). In the gamma V-J pair distribution, a notable pattern emerges with the gene pair TRGV10_TRGJ1 being consistently dominant across all cancer types (Supplementary Figure S5A). Notably, other prevalent gene pairs also involve TRGJ1, including TRGV9_TRGJ1, TRGV2_TRGJ1, TRGV4_TRGJ1, TRGV3_TRGJ1, and TRGV8_TRGJ1. Remarkably, in all cancer types examined, the top 5 most prevalent gene pairs account for over 75% of all γδ-TCR reads, suggesting a shared positive selection mechanism during T cell development. In the delta V-J pair distribution, as illustrated in Figure S5B, TRDV1_TRDJ1 stands out as the dominant gene pair. Other notable pairs include TRDV2_TRDJ1, TRDV1_TRDJ2 and TRDV3_TRDJ1. However, the diversity observed in delta V-J pairs is much lower compared to their gamma counterparts, likely due to the reduced number of reads obtained from the delta chain, which may result from lower expression of the delta chain or the frequent deletion of the TRD locus following TCR gene rearrangement^20^.

### Prognostic implications of **γδ**-TCRs in the pan-cancer dataset

To further explore the prognostic potential of tumor-infiltrating γδ T cells, we performed a systematic survival-analysis screening focused on the gene expression levels of TCR gamma and delta genes across various cancer types (Figure 3). We listed all genes that showed either a significant or marginal impact on patient overall survival, as indicated by their Cox-regression p-values (FDR adjusted) being less than 0.05. In performing the Cox regression analysis, we adjusted for covariates including sex, age, stage, and tumor purity. Please note that in our study, the Cox regression and corresponding p-values were utilized primarily to prioritize the γδ TCR genes for further discussion, rather than to build predictive models or to establish causal relationships. Our analysis identified 24 γδ-TCR genes as potential prognostic biomarkers in 13 cancer types. Among them five genes (*TRGC2*, *TRGV2*, *TRDC,* and *TRDV3*) appeared in more than 3 cancer types, with *TRDC*, often serving as a proxy for the overall presence of γδ T cells^21^, being notably present in 5 different cancers. Another interesting gene, *TRDV3*, a biomarker for the γδ3 subset, has recently attracted attention due to its emerging significance in immune cancer immunology^11^. As expected, the expression of these in cancers like SKCM and HNSC showed an association with favorable survival, contrasting with varied directions in less immunogenic cancer types.

**Figure 3.**
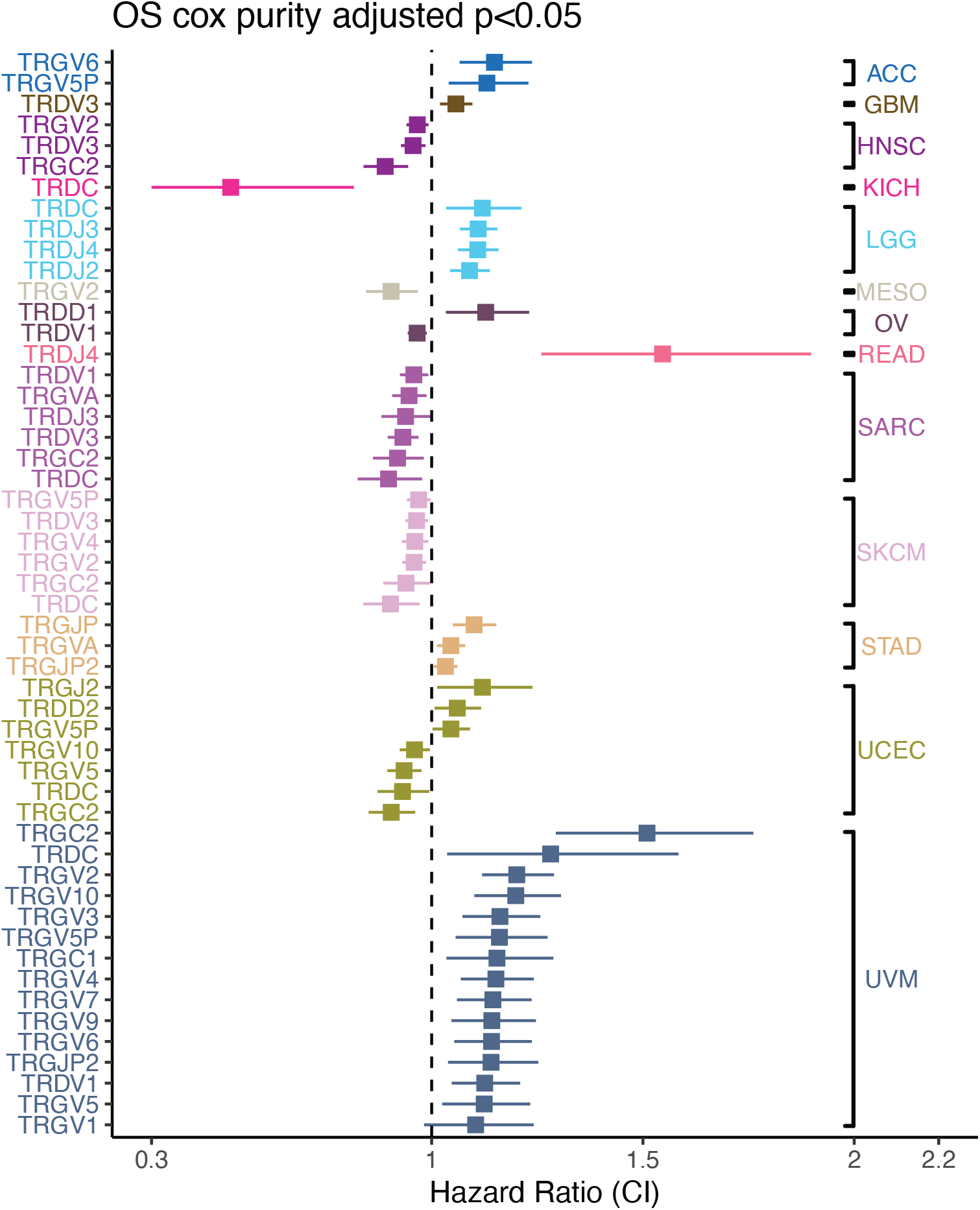
Prognostic implications of gamma-delta TCRs in TCGA. The forest plot displays the results of the Cox regression survival analysis on TCR γ and δ gene expression adjusted by covariates across various cancer types. It highlights genes with significant or marginally significant impacts on patient survival, indicated by FDR-adjusted Cox-regression p-values below 0.05. Each horizontal line in the forest plot represents a γ or δ gene in a cancer type, with the square and length of the line indicating the estimated hazard ratio (HR) and its 95% confidence interval, respectively. Horizontal lines are colored by cancer types. Within each cancer type, genes are ranked by HR from highest to lowest. A dashed vertical line at HR=1 represents no effect.

Acknowledging the complexity inherent in transcriptome-based prognostic signatures, we specifically examined HNSC and COAD, two cancers known for their distinct tumor subtypes (Figure 4). HNSC is categorized into HPV positive and negative subtypes, each with a unique immunological landscape and clinical outcomes. Similarly, COAD exhibits distinct molecular and clinical variability between Microsatellite Instability (MSI) and Microsatellite Stable (MSS) subtypes. Both TCR gamma and delta signatures are notably higher in HPV-positive HNSC patients, suggesting a more active or diverse γδ T cell presence in these patients (Figure 4A). Similarly, the normalized entropy of the TCR gamma signature, a measure indicative of gamma TCR diversity, was also found to be higher in the HPV-positive patient group (Figure 4B). Interestingly, both gamma and delta TCR signatures were significantly associated with patient survival, exhibiting a protective effect in the HPV-positive HNSC subgroup (Figure 4C). However, this association was not observed in the HPV-negative patient group, suggesting the presence of γδ T cells might contribute differently to the disease progress in these subtypes. We also conducted a detailed comparative analysis of HPV-positive and HPV-negative samples, focusing on individual genes, gamma clonality, and genes associated with survival in the stratified groups (Supplementary Figure S6). The analysis indicates that HPV-positive patients typically exhibit a more diverse range of TCR gamma reads. We also observed notable disparities in the prognostic implications of TRG and TRD genes, with these genes appearing more prognostic in the HPV-positive group.

**Figure 4.**
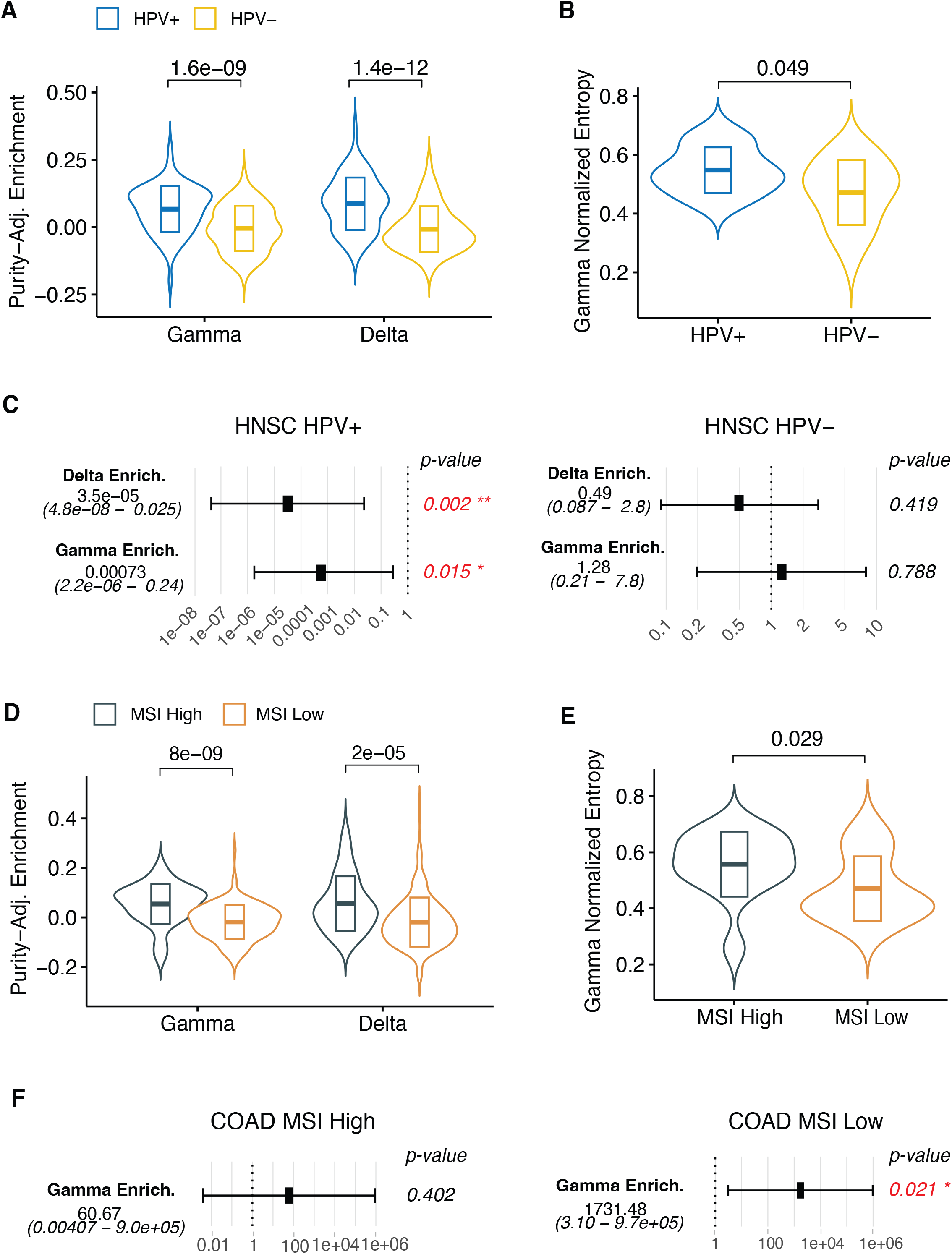
Gamma-delta TCR-based prognostic signatures in TCGA HNSC and COAD. (A) Elevated TCR gamma and delta gene expression in HPV-positive HNSC patients. Y-axis represents the purity-adjusted gamma (left) and delta (right) gene enrichment scores. (B) Higher normalized entropy of TCR gamma signature in the HPV-positive HNSC. Forest plots show the estimated hazard ratio (square) and its 95% confidence interval (horizontal line) of gamma and delta gene enrichment by Cox regression adjusted for covariates in HPV+ and HPV- HNSC, respectively. A dashed vertical line at HR=1 represents no effect. (C) Association of gamma and delta TCR signatures with improved survival in HPV-positive HNSC, absent in HPV-negative. (D) Significantly higher gamma and delta TCR enrichment scores in the MSI-high group. (E) Higher normalized entropy of the gamma TCR in the MSI-high group. (F) The gamma enrichment score associated with worse survival outcomes in the MSI-low group. Forest plots show the estimated hazard ratio (square) and its 95% confidence interval (horizontal line) of gamma gene enrichment by Cox regression adjusted for covariates in MSI high and MSI low COAD tumors, respectively. A dashed vertical line at HR=1 represents no effect. For A-B, and D-E. P-value was calculated by two-sided Wilcoxon rank-sum test. Violin plot shows the kernel probability density of the data with box in middle representing the median (central line), the 25% and 75% interquartile (IQR) (lower and upper hinges).

In the context of COAD, we noted significantly higher values in both gamma and delta TCR enrichment scores in the MSI-high subtype of COAD, alongside an elevation in the normalized entropy of the gamma TCR, indicating a more diverse T cell repertoire in the MSI subgroup (Figures 4D and E). There were differences in the TRG and TRD gene usage and gamma clonality between the two COAD subtypes (Supplementary Figure S7). Intriguingly, we observed that the gamma enrichment score is significantly associated with worse survival outcomes in the MSI-low group of COAD, contrasting with its null association in the MSI-high group (Figure 4F). This disparity reflects the complex and multifaceted role of γδ T cells in influencing cancer prognosis.

### Prognostic implications of **γδ**-TCRs in immunotherapy-treated tumors

We extended our analysis of γδ-TCRs beyond the TCGA pan-cancer dataset by incorporating RNA-seq data from 503 tumors across 8 immunotherapy studies. This additional data allowed us to investigate the prognostic significance of γδ-TCR signatures in the context of immunotherapy. In the Gide study (Figure 5A), we observed that the enrichment of gamma and delta genes was higher in responders (CR/PR) compared to non-responders (SD/PD) in both pre-treatment and on-treatment cohorts, and in the overall patient cohort (Supplementary Figure S8A). Additionally, higher gamma gene enrichment scores were associated with improved survival outcomes in both pre-treatment (middle panel in Figure 5A) and on-treatment samples (Supplementary Figure S8B). Specifically, three individual genes (TRGV2, TRGV8, and TRGV10) demonstrated a protective effect, as shown by the Cox regression analysis of progression-free survival (PFS), adjusted for age and sex. In the Riaz study (Figure 5B), we observed a similar pattern across the entire cohort (left panel) and in the ipilimumab-progressive (Ipi-Prog) patients, but not in the ipilimumab treatment-naïve (Supplementary Figure S8C). Within the Ipi-Prog subgroup (right panel), gamma and delta gene enrichment scores were higher in responders when stratified by pre-treatment and on-treatment tumors. The Kaplan-Meier overall survival plot (Figure 5C) also indicated that patients with higher gamma enrichment scores in pre-treatment samples tended to have better survival outcomes, a similar pattern was observed in on-treatment samples (Supplementary Figure S8D).

**Figure 5.**
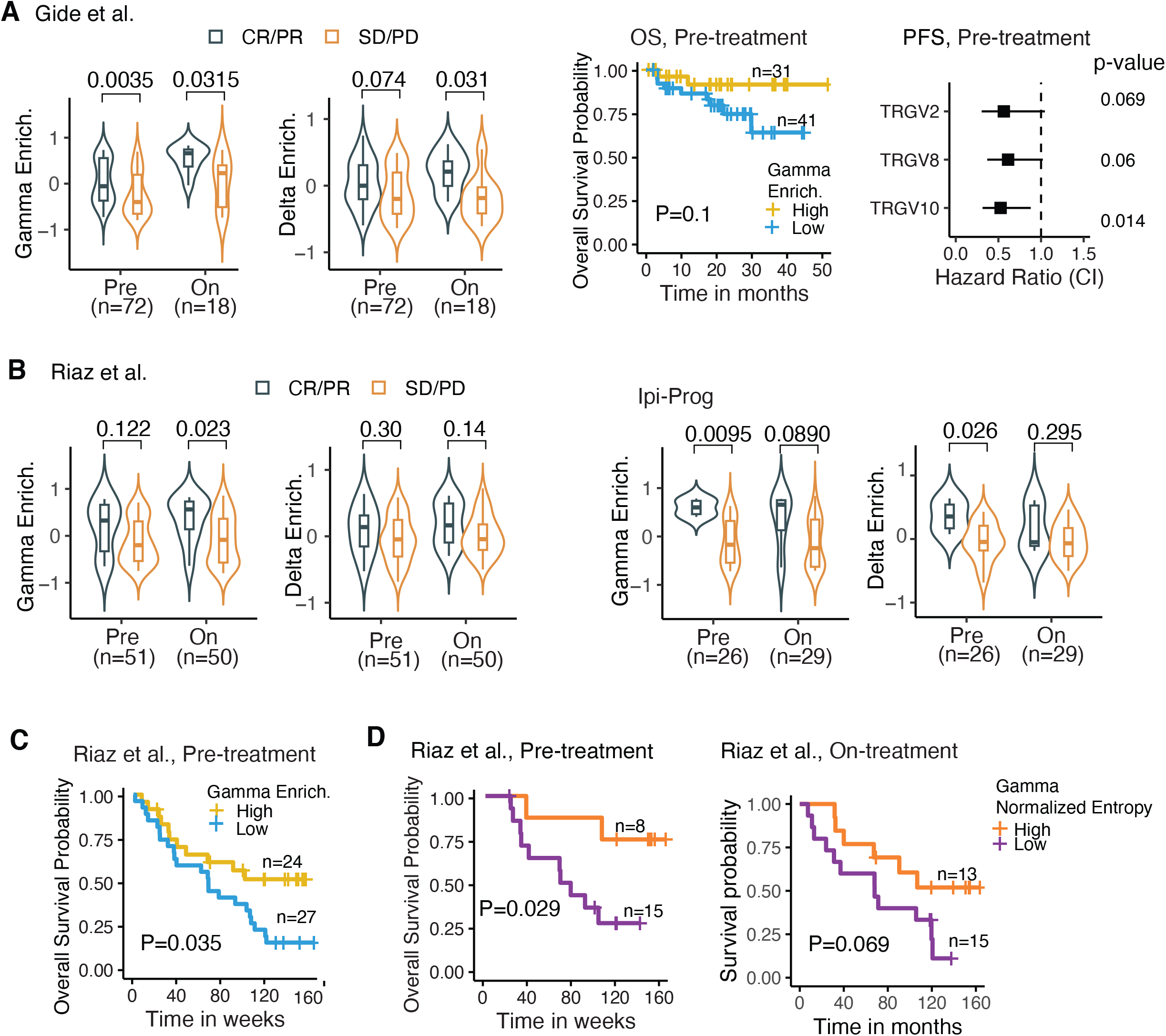
Gamma-delta TCR-based prognostic signatures in immunotherapy studies. (A) Prognostics gamma-delta TCR signatures in Gide study. Violin plots on left panels compare gamma and delta gene enrichment between patients with complete response (CR) or partial response (PR) vs. patients with stable disease (SD) and progressive disease (PD), in pre-treatment and on-treatment samples, respectively. Number of samples in each category was labeled below x-axis. Right panels show the (1) Kaplan-Meier overall survival plot by patients’ pre-treatment samples stratified based on optimal cutoff of gamma gene enrichment, with yellow representing high and blue representing low. (2) Forest plot of PFS (progression free survival) Hazard Ratio (square) and its 95% confidence interval (horizontal line) of gamma gene enrichment by Cox regression adjusted for age and sex. A dashed vertical line at HR=1 represents no effect. P-values are shown on right. (B) Prognostics gamma-delta TCR signatures in Riaz study. Left panel, violin plot with box shows elevated gamma and delta gene enrichment in CP/PD patients vs. SD/PD patients, in both Pre- and On-treatment tumors. Right panel, violin plot with box shows elevated gamma and delta gene enrichment in CP/PD vs. SD/PD, in both Pre- and On-treatment tumors within ipilimumab treatment- progressive (Ipi Prog) patients. (C) In Riaz study, Kaplan-Meier overall survival plot by patients’ pre-treatment samples shows that higher gamma gene enrichment is associated with better survival. Patients are stratified based on optimal cutoff of gamma gene enrichment. (D) Kaplan-Meier overall survival plot by patients’ pre-treatment samples stratified based on optimal cutoff of gamma true diversity in Gide (left) and gamma normalized entropy in Riaz (right), with orange representing high and purple representing low. For violin plots in A-C, P-value was calculated by two-sided Wilcoxon rank-sum test. Violin plot shows the kernel probability density of the data with box in middle representing the median (central line), the 25% and 75% interquartile (IQR) (lower and upper hinges), and the ± 1.5 IQR (Tukey whiskers).

We further analyzed the diversity metrics of gamma chains in immunotherapy studies with patient response information (Supplementary Figure S8E). Our findings indicate that responders, or patients who benefited from treatment, exhibited a more diverse gamma TCR repertoire compared to non-responders across multiple studies. In the Riaz study, we observed that the gamma diversity score, measured by normalized entropy, was associated with improved patient survival in both pre-treatment and on-treatment samples (Figure 5D). A similar pattern was observed in the pre-treatment cohort of the Gide study (Supplementary Figure S8F).

Although based on limited data, these findings confirm the prognostic value of gamma and delta gene expression enrichment and diversity metrics in immunotherapy. Incorporating these γδ-TCR signatures could enhance the precision of patient stratification and treatment personalization when used alongside other established biomarkers.

### Development of an interactive gamma-delta TCR webtool

To facilitate the characterization of gamma-delta TCR repertories, we have developed an open-access resource and web server www.gdt.moffitt.org. This resource curates all gamma-delta repertoires reconstructed with TRUST4, along with gene expression data of gamma-delta TCR genes and clinical information from both TCGA and immunotherapy studies. The web server includes three major modules. (1) *TCR search*. This module allows users to query gamma-delta TCRs identified in both TCGA and immunotherapy studies through CDR3 amino acid sequences (with mismatches allowed) and V/D/J genes. All TCRs matching the querying criteria will be displayed along with their abundance and associated clinical information. (2) *Gene expression analysis*. In this module, users can associate the expression of individual gamma-delta TCR genes with various clinical outcomes through hierarchically clustered heatmaps, in both TCGA and immunotherapy studies. (3) *Prognostic analysis*. Survival analysis (3A): This sub-module provides a comprehensive evaluation of the prognostic significance of features including individual gamma-delta TCR gene expression, gamma/delta gene overall enrichment, and gamma/delta clone diversity. When a feature is queried in a specific cancer type or immunotherapy study, the module renders a Kaplan-Meier plot with either optimal or median cutoff by the choice of users and a forest plot showing the hazard ratio and its confidence interval from a Cox regression adjusted for all covariates. Users can select the desired maximum length of survival time for both Kaplan-Meier and Cox regression analysis. Association with clinical outcomes (3B): This sub-module enables users to explore the gamma-delta gene expression as well as gamma/delta clone diversity with different clinical outcomes, such as responders vs. non-responders, pre- vs. post-treatment, live vs. dead, and various treatment combinations. By providing these functions, this web tool aims to support ongoing research and therapeutic discovery by providing easy access to extensive gamma-delta TCR data and associated clinical insights.

## DISCUSSION

Despite being a minority group in the T cell community, γδ T cells have emerged as a critical area of interest in cancer immunology and translational research. Their dual capability to mirror both innate and adaptive immune responses positions γδ T cells as promising candidates for clinical biomarkers, aiding in the tracking of cancer progression and response to various treatments, including immunotherapy^10,21–24^. Additionally, their complementary immunosurveillance role alongside αβ T cells underscores their potential as novel therapeutic targets in combination therapies^25^. Most of previous studies indicate a protective value for Vδ1+ γδ T cells, but the impact of specific gamma chains that they express and the roles of γδ TILs differing from Vδ1 or Vδ2 on their function remains unclear.

Our comprehensive analysis of the γδ TCR landscape across 33 cancer types, utilizing the TRUST4 algorithm, provides key insights into the role and potential utility of γδ T cell signature in different cancer tissues. To the best of our knowledge, this study represents the largest collection of γδ TCR reads in human cancer to date, providing a critical resource for the ongoing intensive research into this vital immune component. The yielded database offers applications across many research scenarios. For example, users could use it to query the candidate TCR sequences and determine the degree of tumor and tissue specificity. Additionally, they can also compare these sequences against the cancer-specific dataset to identify the common clones or conduct distance-based clustering analysis^26^. Although the variation in CDR3 lengths across cancer types might not provide immediate biological insights, the large collection of the γδ TCR repertoire serves as a valuable reference database. This enables users to detect changes in length distribution (e.g., TCR spectratyping) in individual patients’ TCR repertoires and monitor γδ-specific clonal expansion or selection. Our analysis also highlights the often-overlooked utility of TRG and TRD gene expression values, readily obtainable through standard RNA-seq data analysis. These values offer alternative methods for not only assessing the overall activity of γδ T cells but also for uncovering specific genes that could serve as potential gene-level biomarkers. For instance, in our prior research, we identified *TRGV9* transcripts as key biomarkers in prostate cancer^27^. It is important to acknowledge that expression values of TRG and TRD genes are available in the most current reference annotations for the human genomes, such as GENCODE, but are absent in older versions of the RefSeq gene annotation. Our findings that γδ TCR clones are prevalent across all cancer types, particularly the dominance of specific V-J gene pairs, hinting at a possible link to common antigens or tumor immune response mechanisms. The fact that most immunogenic cancers tend to have higher median TCR γδ gene enrichment scores and clonality further supports the idea of a locally converged immune response in these cancers, possibly against specific tumor antigens.

Recent findings indicate that although in cancer patients, γδ T cell presence correlates with better outcomes, intra-tumor γδ T cells can exhibit a temporal pattern in which their subsets play a different role in supporting anti-tumor immunity^9^. Consequently, it is important to emphasize that any static analysis of γδ TCR clonality or gene expression for prognostic purposes (in testing for the association with patient survival outcomes) should be approached with caution. With this caveat noted, our comprehensive survival analysis has identified multiple TRG and TRD genes as potential prognostic biomarkers across multiple cancer types. Our results in HNSC and COAD also indicate that the role of γδ T cells in disease progression and patient prognosis varies based on the molecular characteristics of the tumor. In HPV-positive HNSC, the higher diversity and clonality of γδ TCRs, and their association with improved survival, contrast with the lack of such associations in HPV-negative HNSC.

Likewise, in COAD, the distinct patterns observed between MSI-high and MSI-low subtypes in terms of γδ TCR enrichment and their association with survival outcomes further emphasize the complex, context-specific contribution of γδ T cells in cancer progression.

Our extended analysis of γδ-TCRs in immunotherapy datasets, including RNA-seq data from 503 tumors across 8 studies, underscores the significant role of γδ T cells in predicting patient response to immunotherapy. In multiple studies, we observed that patients with immunotherapy benefits tend to exhibit higher gamma and delta gene enrichment scores, as well as higher diversity in gamma chain TCRs. These results suggest that incorporating γδ-TCR signatures could further enhance the precision of treatment personalization in immunotherapy. Although our findings highlight the prognostic value of intra-tumor γδ T cells and specific TRG and TRD genes, we must acknowledge that the true therapeutic implications of γδ T cells are multifaceted. Different γδ T cell subsets are activated by distinct stimuli, such as BTNL3 and BTNL8 for Vγ4Vδ1 cells ^28,29^, which suggests that tailored approaches to activate these cells could enhance their anti-tumor effects. Therapeutic strategies currently under investigation include reactivating Vγ9Vδ2 T cells with specific antibodies and the adoptive transfer of γδ T cell subsets. Current trials using Vδ1 and Vδ2 T cells have shown promise in treating solid tumors, hematological malignancies, and bone marrow metastases ^30–32^.

Understanding which γδ T cell subsets naturally home to and persist within tumor beds is critical for developing effective cellular therapies. The database generated by our study provides a valuable resource for advancing these therapeutic strategies, offering new avenues for personalized immunotherapy that harness the unique properties of γδ T cells to improve patient outcomes.

A major analytical challenge of this study arises from the fact that γδ T cells constitute only a small proportion of lymphocytes, which can complicate their detection and analysis with bulk tumor RNA-seq data. This issue is particularly problematic in cancers with lower immune infiltration, where the presence of γδ T cells may be even more sparse. Furthermore, the delta chain analysis encounters additional hurdles due to its lesser representation and fewer annotations or references available compared to the gamma chain. Consequently, most of our analysis focused on the Gamma TCR. Even in tumors with higher levels of γδ T cells, it remains challenging to differentiate the prognostic signature from overall inflammation or immune infiltration, similar to other immune-related signatures. To address this issue, we performed a sensitivity analysis by adjusting for the overall immune infiltration score (IIS) in the Cox regression model (Supplementary Figure S9). Encouragingly, most of the significant genes retained their significance, indicating their prognostic value is at least partially independent of overall immune infiltration.

Another intriguing question warranting further investigation is whether γδ T cells are more represented in the tumor microenvironment compared to normal tissues. This analysis is challenging with existing data because many normal tissues in cancer genomic datasets are adjacent normal tissues, representing a unique compartment. It is known that γδ T cells tend to patrol tissues for signs of infection or cellular stress, therefore they might be enriched in regions closer to tumor. Additionally, normal tissue RNAseq often receives lower sequencing depth, resulting in a lower detection rate of γδ T cell-related signatures. Despite these limitations, for the sake of completeness, we analyzed all cancer types with at least five samples with matched-normal-tissue RNA-seq and found that the diversity of gamma TCRs is significantly higher in breast cancer and lung squamous cell carcinoma (Supplementary Figure S10).

This atlas will continue to expand by incorporating more TCR repertoires from published studies, including data from both bulk RNA-seq and single-cell analyses, along with additional functional annotations. Because γδ T cells are tissue-specific and functionally plastic, single-cell RNA-seq would be particularly useful for further investigating their functional roles, cytotoxicity, and effector subsets.

Emerging data from tissue-derived γδ T cells ^11,33,34^, obtained from both animal models and human studies, highlight the importance of such detailed analyses to better understand the diverse roles these cells play in the immune response and their interactions within the tumor microenvironment. This expansion will enable users to efficiently integrate their TCR data with the most comprehensive and up-to-date tumor-specific repertoire available, facilitating a more comprehensive view of the γδ-T cell-related immune landscape in cancer and potentially uncovering novel therapeutic avenues.

## METHODS

### Data download and processing

Pre-processed RNA-seq BAM files of 10,970 TCGA samples generated by STAR^35^ were downloaded from Genomic Data Commons (GDC) through controlled data access application from dbGAP study accession phs00178. Gene counts and Fragments Per Kilobase of transcript per Million (FPKM) expression values (Gencode V36) of TCGA tumors generated by STAR were also obtained from GDC. According to GDC, the raw sequencing reads were first accessed for quality using fastqc and then aligned to GRCh38 with Gencove V36 using STAR two-pass mode, which includes a splice junction detection step. Post-alignment quality assessment was conducted using Picard. Only samples pass the quality criteria of GDC were released. FPKM values of gamma-delta TCR genes we extracted from GDC and used for this study are hosted at www.gdt.moffitt.org. Clinical information including overall survival, age, sex, stage, tumor purity of TCGA patients was downloaded from PanCancerAtlas ^36^ (TCGA-CDR-SupplementalTableS1.xlsx). A consensus measurement of purity estimations (CPE) ^37^ derived from purity level ABSOLUTE, ESTIMATE, LUMP, and IHC was used in downstream analysis. For samples without CPE value available, purity estimated by ABSOLUTE downloaded from PanCancerAtlas (TCGA_mastercalls.abs_tables_JSedit.fixed.txt) were used to represent tumor purity. Microsatellite instability (MSI) scores of COAD and HPV status of HNSC were obtained from cBioPortal. The MSI scores were represented by MANTIS ^38^, which was the average values of MSI after calculating the allele distribution differences of each microsatellite site between tumor and normal samples. COAD tumors were further categorized into MSI-high vs. MSI-low groups, with the cutoff determined by identifying the local minimum in the bimodal distribution of MSI values density plot (Supplementary Figure S7A). For HNSC tumors without HPV status available on cBioPortal, we imputed their status based on K-means clustering analysis using the top 100 differentially expressed genes between HPV+ and HPV- tumors.

### TCR reconstruction

The RNA-seq BAM files downloaded GDC from were processed by TRUST4 v1.0.4 to reconstruct immune receptor repertories in T cells along with their abundances using command line run-trust4 -f human_IMGT+C.fa --ref human_IMGT+C.fa -b sample.bam -o sample_trust4. TRUST4 reported complete immune receptor CDR3 sequences, where the sequence started from the IMGT-specified (https://www.imgt.org/) V gene location and ended with the IMGT-specified J gene location. The abundance for each CDR3 in this study was estimated using the 3 million candidate TCR reads overlapping CDR3 regions when executing TRUST4. A total of 34,129 CDR3 sequences of gamma-delta (γδ) T cells from 6,751 tumors of 6,392 patients were subjected for analysis. We applied following filtering criteria to remove false positive calls and non-informative TCRs:

1. CDR3 sequences from patients lacking associated clinical information were excluded.
2. For patients with multiple tumor samples, only the CDR3 sequences assembled form the primary tumor sample with the highest number TCRs identified were retained for the analysis.
3. CDR3 sequences carrying alpha-beta V genes were filtered out.
4. Out-of-frame CDR3 sequences (459 γ, 8,920 δ) were removed.

To create unique identifiers of γδ-TCR clones, we used a combination of CDR3 amino acid sequences, V genes, and J genes. Finally, 22,205 distinct γδ-TCR clones (4,554 γ clones, 17,651 δ clones) from 6,276 tumors across 33 cancer types were retained for repertoire analysis.

### Gamma-delta TCR repertoire analysis

Following metrics were calculated to the explore repertoire of gamma and delta TCR clones.

***Normalized chain count***: For each tumor, gamma (or delta) chain count was calculated as log 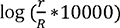, where *r* is the total count of reads mapped to any gamma (or delta) clones, and *R* is the total number of RNA-seq reads mapped to transcriptome as in STAR count data downloaded from GDC. ***Clone size***: Clone size was defined as total number of reads mapped to each clone. Clone size distribution was analyzed across 33 different cancer types by categorizing clones into sizes of 1, 2, 3- 10, 11-100, or >100 reads visualized by vinplot function in *ggplot2*. ***V-J gene usage***: The usage of V-J gene pairs was explored by calculating the number of distinct clones (counts) and percentage of CDR3 reads (frequency) harboring a specific combination of V-J pair. V-J pairs with a frequency higher than 0.3% were visualized in stacked bar plots for each cancer type using *ggplot2*. ***CDR3 length***: CDR3 amino acid length distribution analysis was conducted across 33 cancer types in gamma and delta clones respectively and visualized in ridge plot in *ggplot2*. Cancer types were ranked by the mode (max values) of CDR3 length density distribution. ***Clonality and entropy***: For each tumor with at least 5 distinct clones, the clonality of gamma and delta clones was measured by Shannon clonality 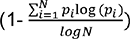 and Simpson clonality 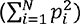, where *p_i_* is the frequency of clone *i*, and N is the total number of distinct clones. We also calculated normalized Shannon entropy as 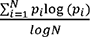 to measure diversity. Clonality close to 0 characterizes a polyclonal repertoire with equivalent representations of each clone (*i.e.* infinite diversity); clonality close to 1 represents a highly oligoclonal repertoire (*i.e.* no diversity). We compared the clonality and entropy of gamma-delta clones between HPV-positive and HPV-negative HNSC tumors and between MSI-high and MSI-low COAD tumors using the Wilcoxon signed-rank test.

### Gamma-delta TCR gene expression

The overall enrichment of gamma (or delta) gene expression was calculated from TPM values downloaded from GDC using single-sample GSEA implemented in R package *GSVA* on all TCR gamma (or delta) genes with default setting. In order to correlate normalized gamma (or delta) TCR clone abundance with gamma (or delta) gene expression enrichment scores in each cancer type, partial Spearman rank correlation controlling for tumor purity of each tumor was performed using pcore.test function in *ppcor* R package with method=”spearman”. The prognostic potentials of individual gamma delta TCR genes (V, D, J, C) expression were evaluated by Cox proportional hazard regression analysis on TPM values adjusted for covariates including sex, age, stage, and tumor purity across different cancer types, using coxph function of R package *survival*. *P* values were further adjusted by the Benjamini–Hochberg method to correct for multiple comparisons. The Hazard Ratio and confidence interval were visualized for gamma-delta TCR genes with adjusted *p*-value < 0.05. We also examined the correlation of the expression of individual gamma delta TCR genes and TCR enrichment scores with different tumor subtypes such as: HPV-positive vs. HPV-negative HNSC tumors, and MSI-high vs. MSI-low COAD tumors. Specifically, TCR gene TPM values and enrichment scores were first adjusted for tumor purity by taking residuals from linear regression; then Wilcoxon signed-rank test was performed to compare the purity adjusted values between HPV-positive vs. HPV-negative HNSC tumors, as well as between MSI-high vs. MSI-low COAD tumors; results were visualized by boxplot function in *ggplot2*. In addition, to assess differences in the prognostics potentials of expression levels of individual TCR genes among above different tumor subtypes, we conducted Cox proportional hazard regression adjusting for sex, age, stage, and tumor purity using coxph function of R package *survival*. *P* values were further adjusted by the Benjamini–Hochberg method. The Hazard Ratio (and confidence interval) and adjusted *p*-values were visualized for genes showing different prognostics values between different tumor subtypes for HNSC and COAD. Similarly, cox regression adjusted for sex, age, stage, and tumor purity was performed and visualized to evaluate the different prognostics potentials of TCR enrichment scores between HPV-positive vs. HPV-negative HNSC tumors, as well as between MSI-high vs. MSI-low COAD tumors.

### Data analysis of Immunotherapy studies

RNA-seq reads of eight immunotherapy studies were downloaded from GSE78220 (Hugo et al.^39^), GSE91061 (Riaz et al. ^39^), GSE121810 (Cloughesy et al.^40^), GSE165252 (van den Ende et al.^41^), GSE183924 (Mamdani et al.^42^), PRJEB23709 (Gide et al.^43^), PRJEB25780 (Kim et al.^44^), and PRJNA923698 (Campbell et al.^45^). The raw sequencing reads were first assessed for quality using FastQC (http://www.bioinformatics.babraham.ac.uk/projects/fastqc/). Quality trimming were performed using cutadapt ^46^ to remove reads with adaptor contaminants and low-quality bases. Reads pairs with either end too short (<25bps) were discarded from further analysis. Strandness of reads was inferred by RSeQC infer_experiment.py ^47^ with default setting. Next, trimmed and filtered reads were aligned to the human transcriptome GRCh38 using STAR ^35^ with parameters --outSAMtype BAM SortedByCoordinate --readFilesCommand zcat --twopassMode Basic --chimOutType Junctions SeparateSAMold --outFilterMismatchNoverLmax 0.04 --chimSegmentMin 10 --quantMode TranscriptomeSAM. Gene-level TPM (transcripts per million) values were obtained by RSEM ^48^ using Gencode V30 with parameters --star --star-gzipped-read-file --estimate-rspd --strandedness S, where S is strandness inferred by RSeQC. Overall enrichment of gamma (or delta) gene expression was then calculated from TPM values using ssGSEA as described above. Wilcoxon signed-rank test was performed to compare gamma (or delta) enrichment between responders vs. non-responders, and pre-treatment vs. after-treatment. For Riaz study, we split the patients into ipilimumab treatment naïve (Ipi Naïve) and progressive (Ipi Prog) following the original paper. For studies with survival data available, Kaplan-Meier survival analysis was performed to compare overall survival between groups using optimal gamma (or delta) enrichment as cutoff. The optimal cutoff was determined using surv_cutpoint function in R package surminer. The prognostic potentials of individual gamma delta TCR genes (V, D, J, C) expression were evaluated by Cox proportional hazard regression analysis on TPM values adjusted for sex and age using coxph function of R package survival.

Immune receptor repertories of T cells were constructed from the raw sequencing reads using TRUST4 v1.0.4 with command line as following. Paired-end data: run-trust4 -f human_IMGT+C.fa -- ref human_IMGT+C.fa -1 sample_1.fastq -2 sample_2.fasta -o sample_trust4; Single-end data: run- trust4 -f human_IMGT+C.fa --ref human_IMGT+C.fa -1 sample.fastq -o sample_trust4. CDR3 sequences and γδ-TCR clones were identified and filtered as described above in TCR reconstruction section for TCGA. A total of 3,574 γδ-TCR clones (2,887 γ clones, 687 δ clones) from 469 tumors across 8 immunotherapy studies were retained for repertoire analysis. Three metrics were calculated to measure diversity of gamma-delta TCR repertoire: True diversity (effective number of species), 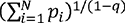; Normalized Shannon entropy, 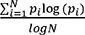; Inverse Simpson index, 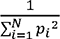, where *p_i_* is the frequency of clone *i*, N is the total number of distinct clones, and *q* = 5. Wilcoxon signed-rank test was performed to compare gamma (or delta) diversity between responders vs. non-responders, and pre-treatment vs. after-treatment. Kaplan-Meier survival analysis was performed to compare overall survival between groups with high vs. low true diversity, with optimal cutoff determined using surv_cutpoint function in R package surminer.

## Data Availability

RNAseq BAM files and FPKM gene expression values were downloaded from Genomic Data Commons (GDC). Clinical information, FPKM values of all gamma-delta related genes, and diversity measurements are available at www.gdt.moffitt.org. Raw RNA-seq data of eight Immunotherapy studies were obtained from Gene Expression Omnibus (GEO) repository with following accession numbers: GSE78220 (Hugo et al.^39^), GSE91061 (Riaz et al. ^39^), GSE121810 (Cloughesy et al.^40^), GSE165252 (van den Ende et al.^41^), GSE183924 (Mamdani et al.^42^), and from European Nucleotide Archive (ENA) with following number: PRJEB23709 (Gide et al.^43^), PRJEB25780 (Kim et al.^44^), and PRJNA923698 (Campbell et al.^45^). Clinical information, processed gene expression in TPM values, and diversity measurements of gamma-delta TCRs are available at www.gdt.moffitt.org.

## ACKNOWLEDGEMENTS

This work has been supported in part by a National Institute of Health grant, R01DE030493 and P20GM130454 (Dartmouth); and Moffitt’s Biostatistics and Bioinformatics Shared Resources at the H. Lee Moffitt Cancer Center & Research Institute, an NCI-designated Comprehensive Cancer Center (P30-CA076292).

