## Supplementary Figures S1-S10 for "A pan-cancer gamma delta T cell repertoire"

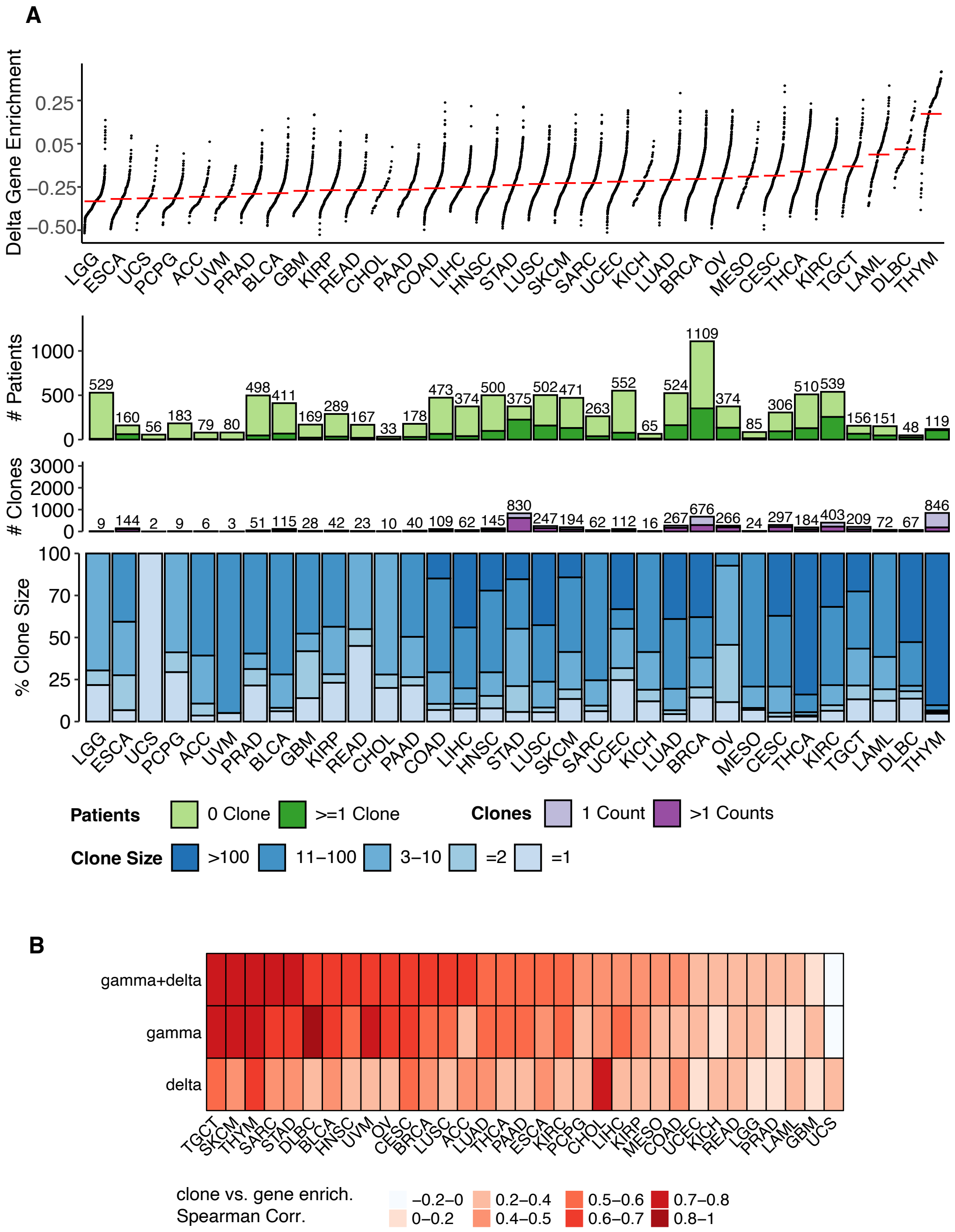


**Supplementary Figure S1.** Landscape of TCR delta gene expression and clones across cancer types. (A) Variability in TCR delta gene enrichment and clonotype presence across 33 distinct TCGA cancer types. Top panel shows delta gene enrichment scores of tumors. Each dot represents a tumor, with y-axis position indicating the normalized delta gene enrichment score. Cancer types are ordered by their median enrichment scores, with the lowest scores on left and highest on right. The second panel shows the number of patients in each cancer types, with light green indicating patients without any δ TCR identified and dark green indicating patients with at least one δ TCR clone identified. The third panel shows number of unique δ TCR clones identified in each cancer type, with light purple representing clones with only 1 read and dark purple representing clones with > 1 reads. The bottom panel shows the proportion of unique δ TCR clones with specific clone sizes. (C) Spearman correlation between γ, δ, and γ+ δ gene enrichment scores vs. normalized chain counts within each cancer type.

**Gamma Chains**

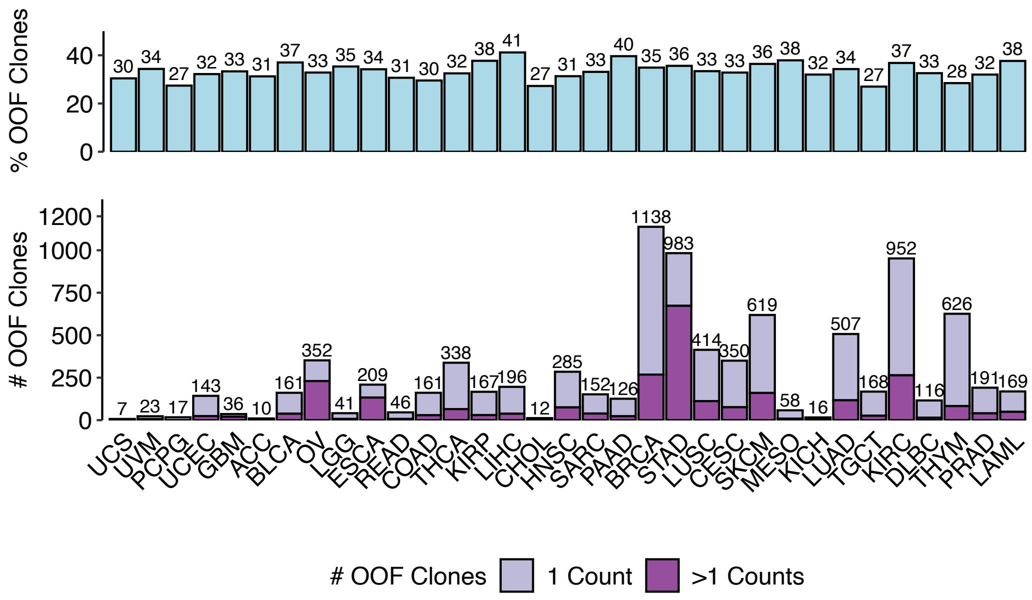


**Delta Chains**

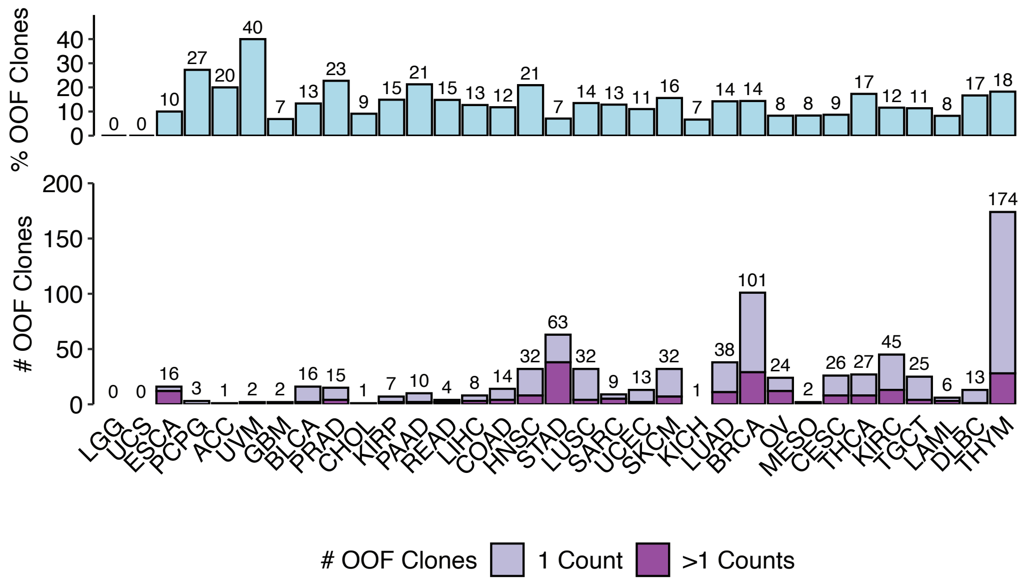


**Supplementary Figure S2.** Distribution of out-of-frame TCR across cancer types in TCGA. (A) Gamma chains. Top panel shows percentage of out-of-frame (OOF) gamma clones. Bottom panel shows number of OOF gamma clones identified in each cancer type. (B) Delta chains. Top panel shows percentage of out-of-frame (OOF) delta clones. Bottom panel shows number of OOF delta clones identified in each cancer.

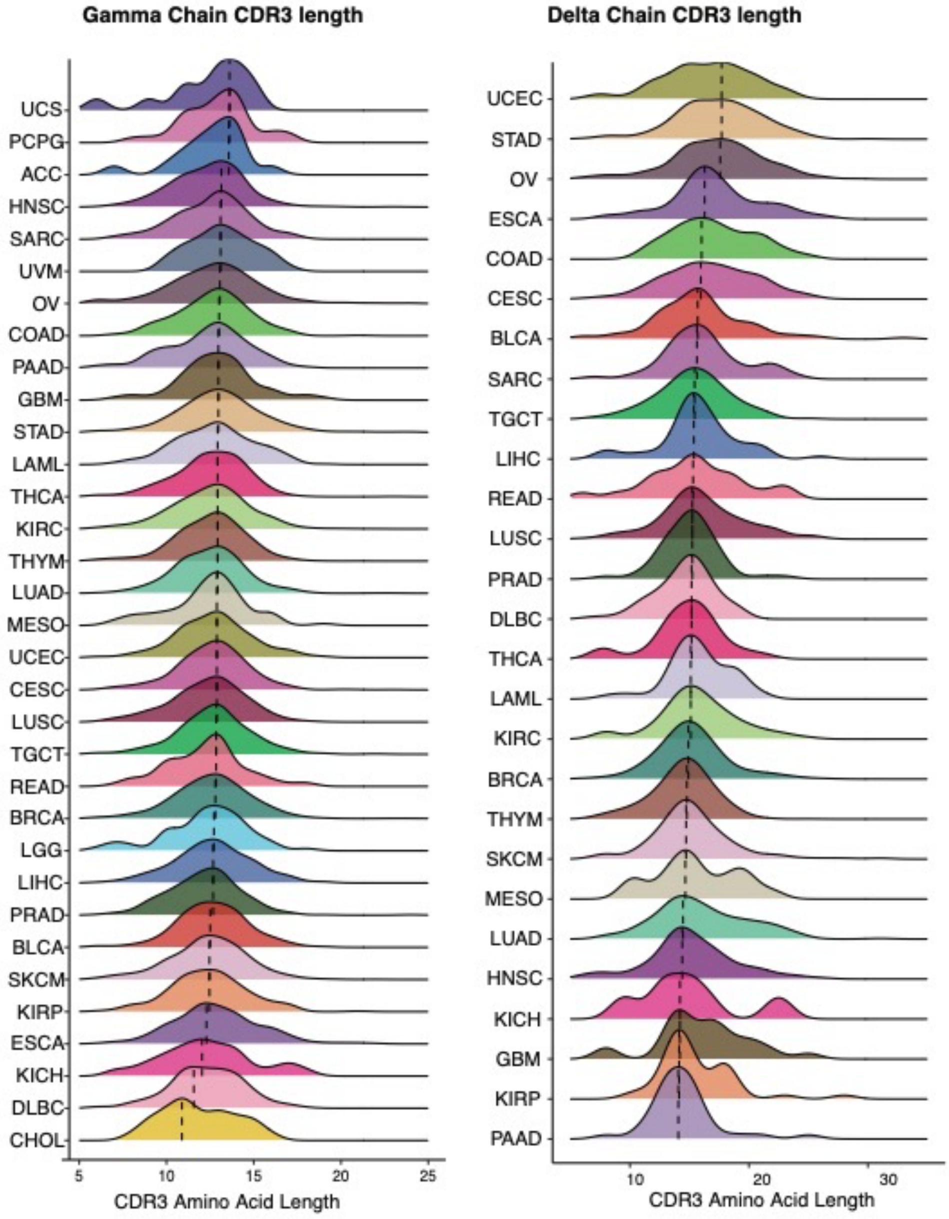


**Supplementary Figure S3.** CDR3 length distribution in gamma (left) and delta (right) TCR across TCGA cancer types. Cancer types are ranked by mode (max values) of CDR3 length density distribution.

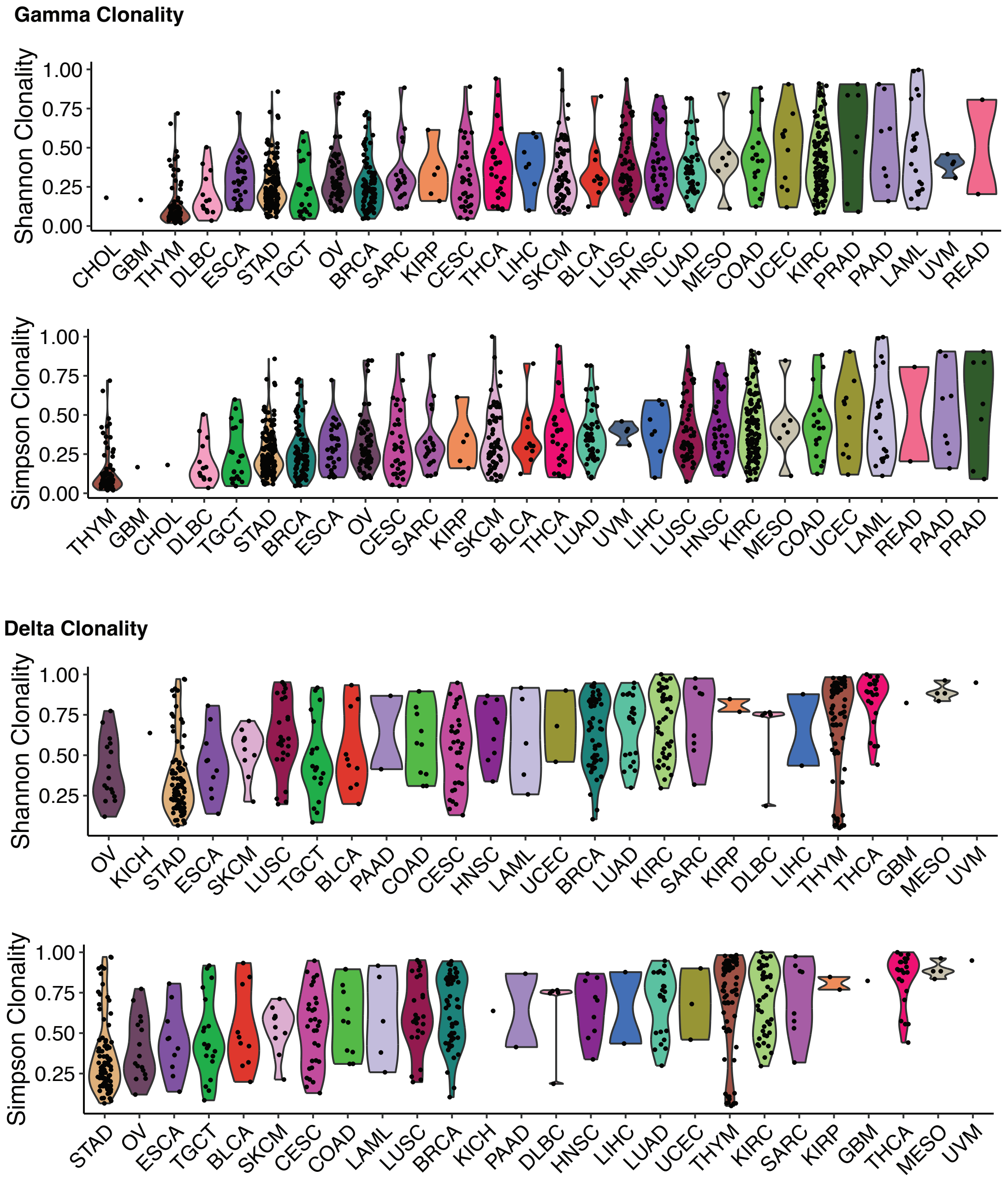


**Supplementary Figure S4.** Shannon and Simpson Clonality values across cancer types in TGCA.

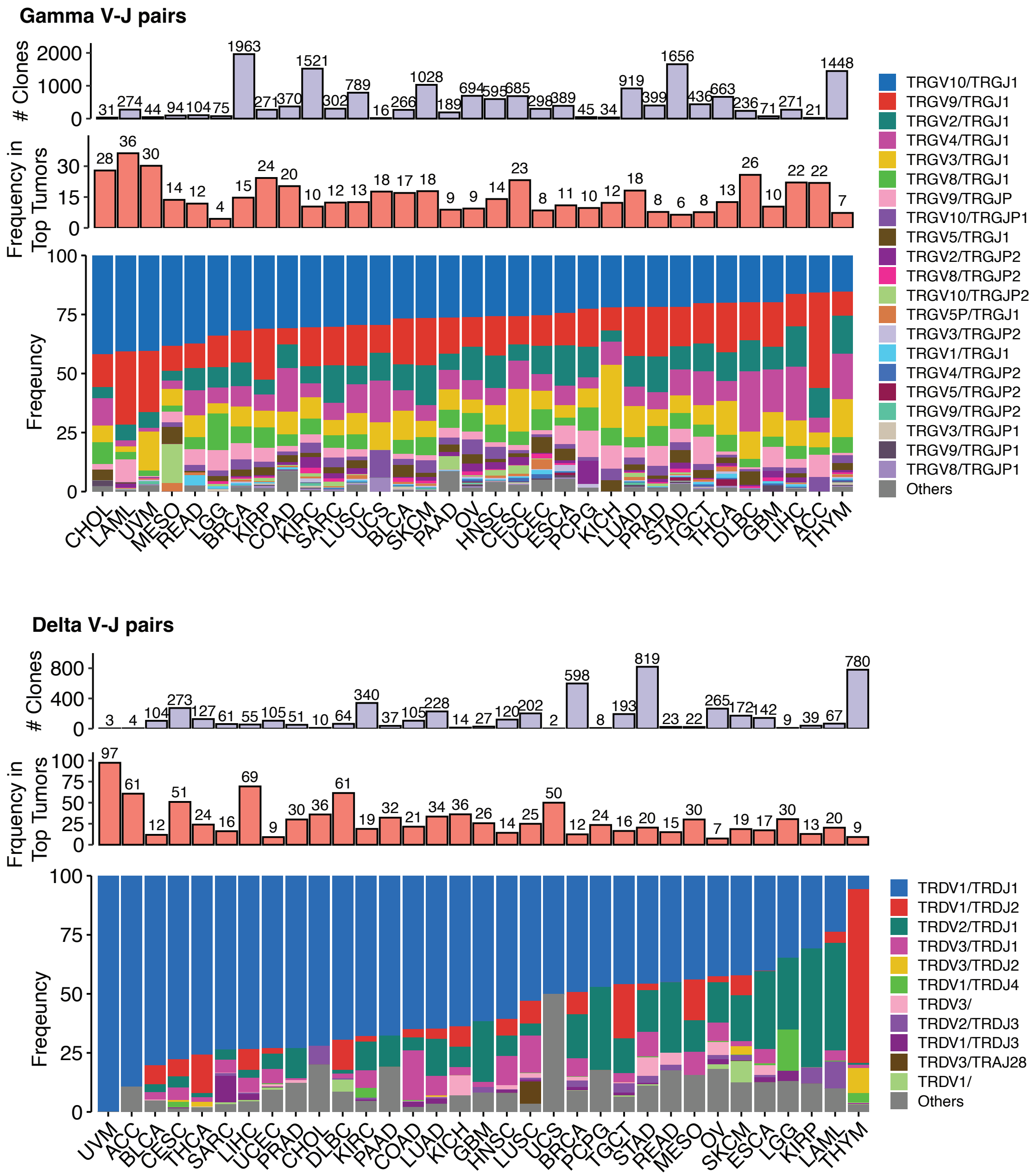


**Supplementary Figure S5.** V-J gene pair distribution on gamma (top) and delta (bottom) TCRs. Top, three panels from top to bottom show the number of gamma clones, percentage of gamma clones in top 10% tumors with the highest gamma chain counts, frequency of different gamma V-J gene pair usage across cancer types. Cancers are ranked by frequency of TRGV10/TRGJ1 from highest to lowest. Bottom, three panels from top to bottom show the number of delta clones, percentage of delta clones in top 10% tumors with the highest delta chain counts, frequency of different delta V-J gene pair usage across cancer types. Cancers are ranked by frequency of TRDV1/TRDJ1 from highest to lowest.

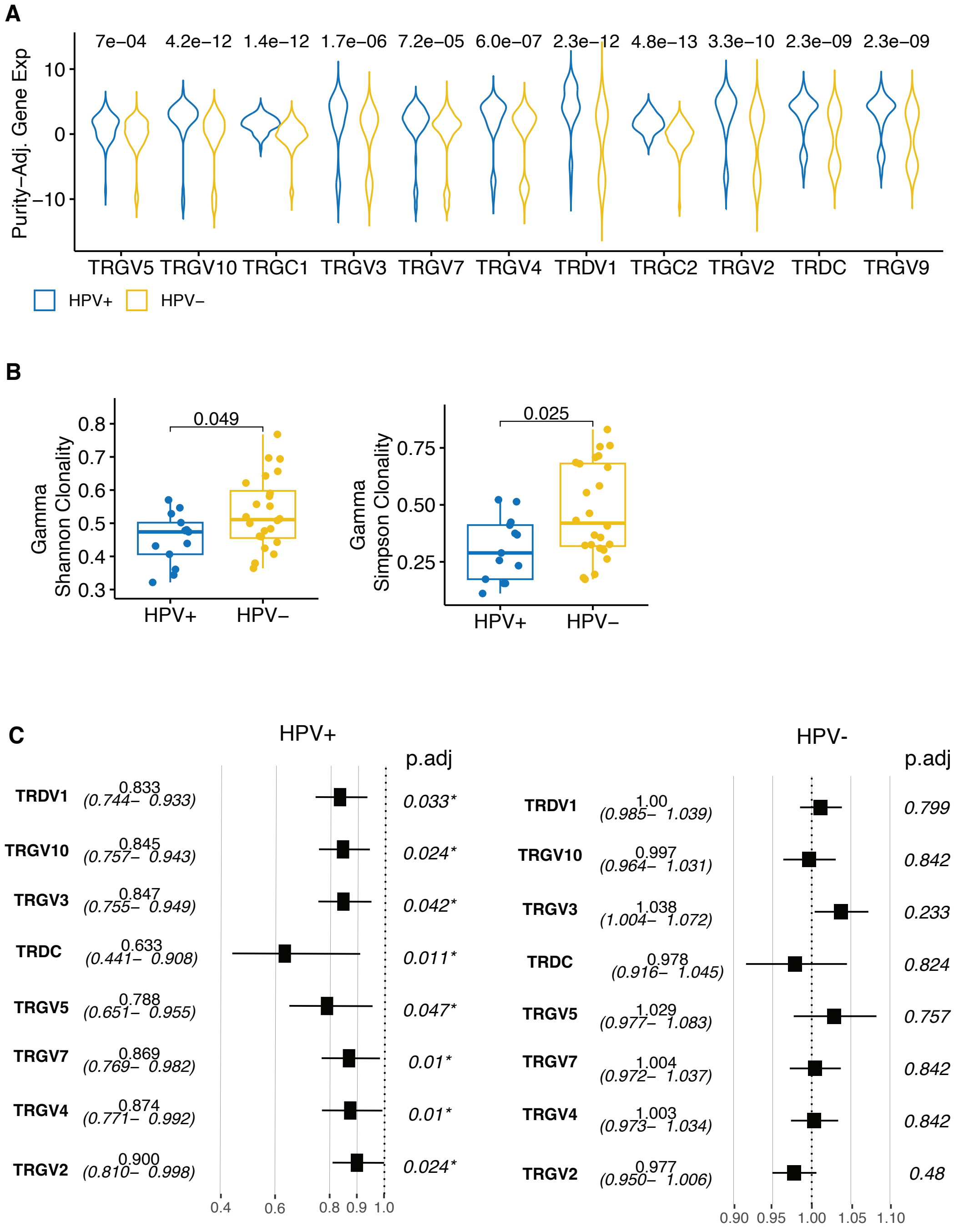


**Supplementary Figure S6.** Comparative analysis of TCR signatures in HPC+ and HPV- HNSC cancer in TCGA. (A) Violin plot of purity-adjusted gamma gene expression between HPV+ vs. HPV-. P-value was calculated by two-sided Wilcoxon rank-sum test. (B) Boxplot comparing gamma Shannon clonality (left) and Simpson clonality (right) between HPV+ vs. HPV-. P-value was calculated by two-sided Wilcoxon rank-sum test. Box represents the median (central line), the 25% and 75% interquartile (IQR) (lower and upper hinges), the ± 1.5 IQR (Tukey whiskers), and all data points.
. (C) Forest plot of overall survival Hazard Ratio (square) and its 95% confidence interval (horizontal line) of individual gamma and delta gene by Cox regression adjusted for covariates for HPV+ (left) and HPV- (right). A dashed vertical line at HR=1 represents no effect. FDR adjusted p-values are shown on right of each panel. Estimated Hazard Ratio and 95% confidence interval are shown beside gene names.

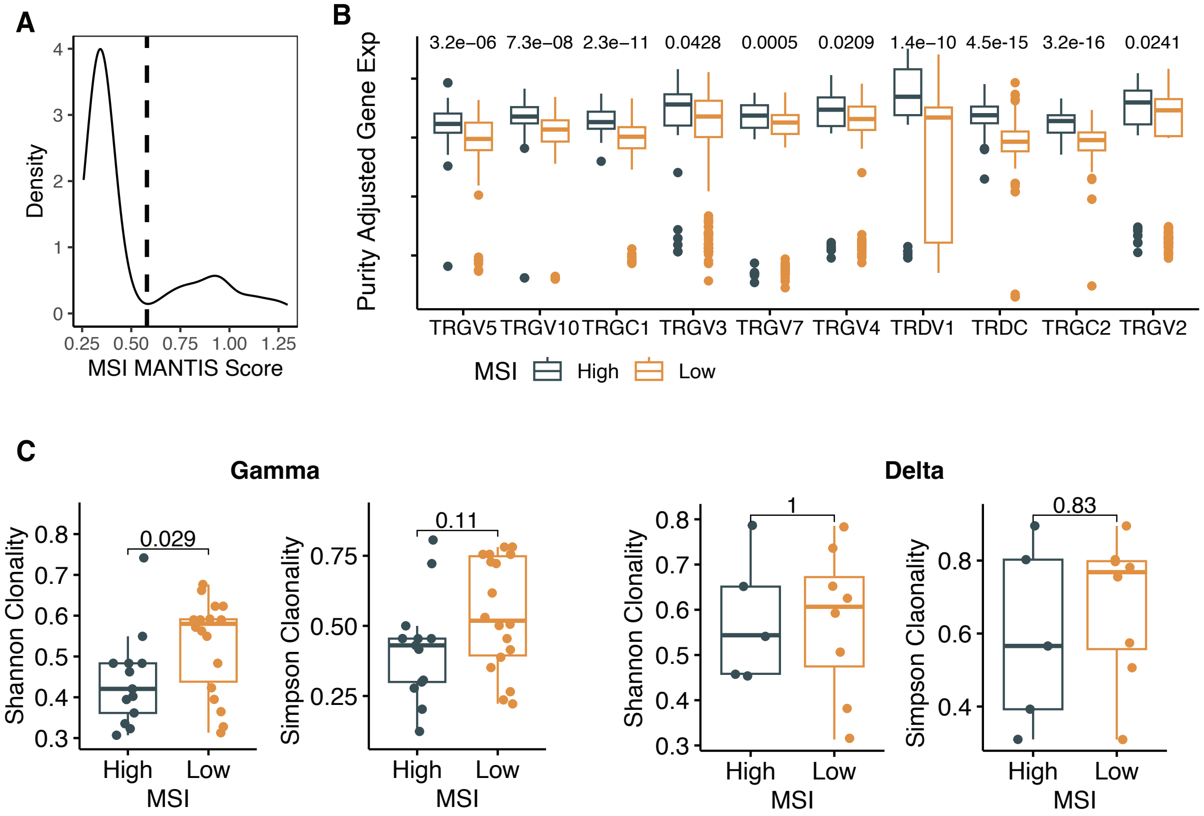


**Supplementary Figure S7.** TCR signature and clonality in COAD subtypes. (A) Distribution of MSO MANTIS score in COAD cohort. The dashed line represents the cutoff determined by identifying the local minimum in the bimodal distribution. Tumors on left of the dashed line are defined as MSI-low. Tumors on right of the dashed line are defined as MSI-high. (B) Boxplot showing gamma and delta TCR gene usage difference in MSI-high and MSI-low group. Y-axis denotes the purity-adjusted gene expression value. P-value was calculated by two-sided Wilcoxon rank-sum test and adjusted by FDR. (C) Left panel, boxplot comparing gamma Shannon clonality and Simpson clonality between MSI-high vs. MSI-low COAD tumors. Right panel, boxplot comparing delta Shannon clonality and Simpson clonality between MSI-high vs. MSI-low COAD tumors. P-value was calculated by two-sided Wilcoxon rank-sum test. Box represents the median (central line), the 25% and 75% interquartile (IQR) (lower and upper hinges), the ± 1.5 IQR (Tukey whiskers), and all data points.

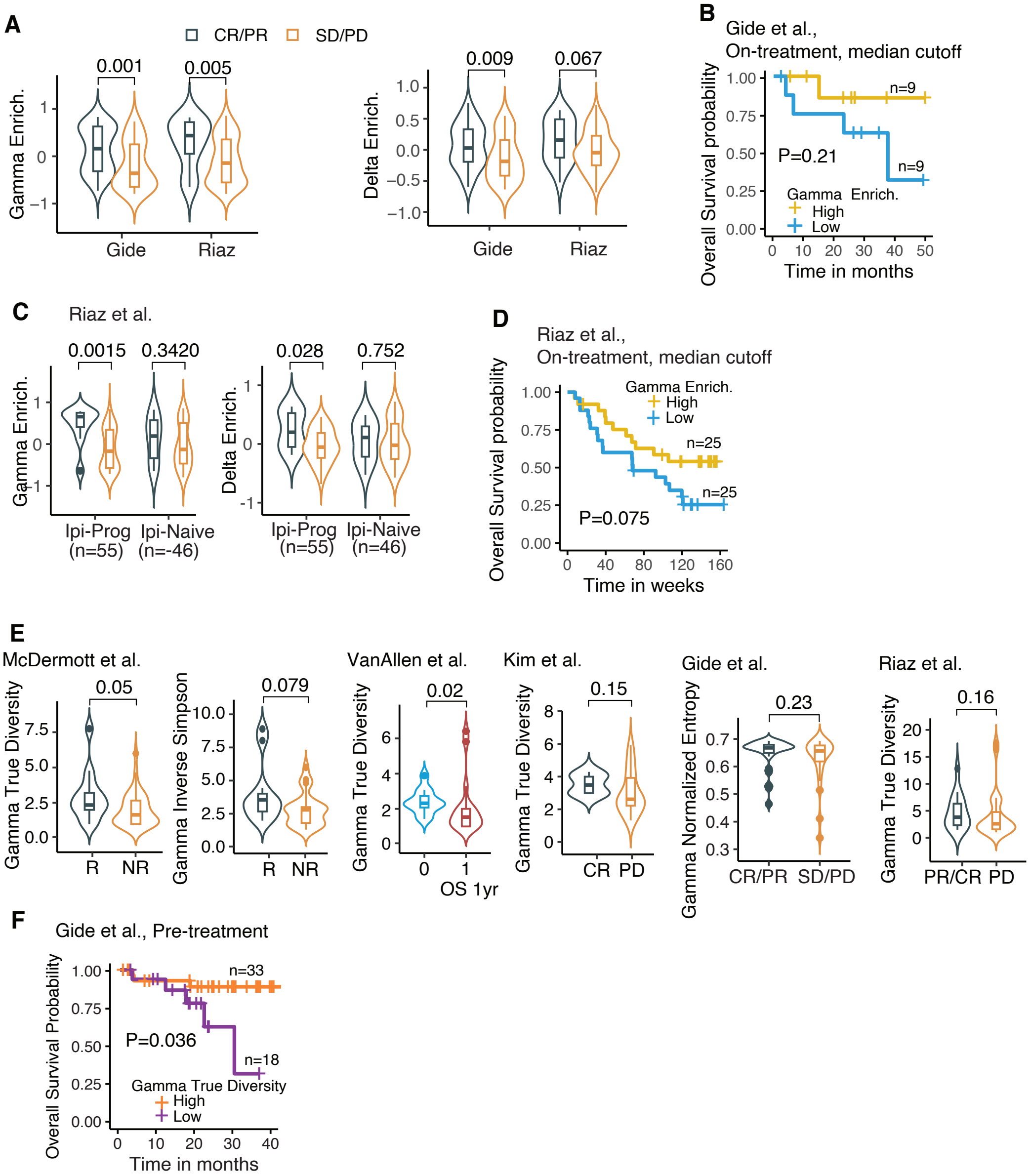


**Supplementary Figure S8.** Prognostics analysis of gamma-delta TCR related features in immunotherapy studies. (A) Violin plots compare gamma (left) and delta (right) gene enrichment between patients with complete response (CR) or partial response (PR) vs. patients with stable disease (SD) and progressive disease (PD), in Gide and Riaz studies. (B) In Gide study, Kaplan-Meier overall survival plot by patients' pre-treatment samples stratified based on median cutoff of gamma gene enrichment, with yellow representing high and blue representing low. (C) In Riaz, violin plot with box shows elevated gamma and delta gene enrichment in CP/PD patients vs. SD/PD patients, in both ipilimumab treatment naïve (Ipi Naïve) and progressive (Ipi Prog) patients. (D) In Riaz, Kaplan-Meier overall survival plot by patients' pre-treatment samples stratified based on median cutoff of gamma gene enrichment, with yellow representing high and blue representing low. (E) Violin plot of gamma true diversity, gamma inverse simpson index between responders (R) vs. non-responders (NR) in McDermott, Kim, Riaz, and Gide studies. Violin plot of gamma true diversity in VanAllen study between patients did vs did not survive > 1 year. (F) Kaplan-Meier overall survival plot by patients' on-treatment samples stratified based on optimal cutoff of gamma normalized entropy in Gide, with orange representing high and purple representing low. For violin plots in A and D, P-value was calculated by two-sided Wilcoxon rank-sum test. Violin plot shows the kernel probability density of the data with box in middle representing the median (central line), the 25% and 75% interquartile (IQR) (lower and upper hinges), and the ± 1.5 IQR (Tukey whiskers).

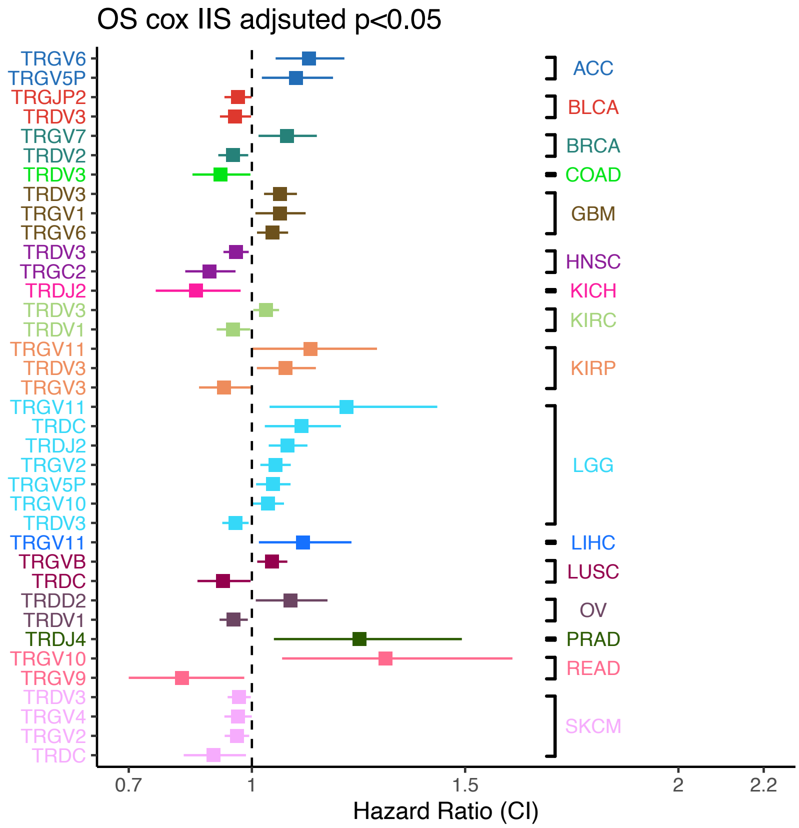


**Supplementary Figure S9.** Cox regression on TCR γ and δ gene expression adjusted by immune infiltration scores (IIS) and other covariates. It highlights genes with significant or marginally significant impacts on patient survival, indicated by FDR-adjusted Cox-regression p-values below 0.05. Each horizontal line in the forest plot represents a γ or δ gene in a cancer type, with the square and length of the line indicating the estimated hazard ratio (HR) and its 95% confidence interval, respectively. Horizontal lines are colored by cancer types. Within each cancer type, genes are ranked by HR from highest to lowest. A dashed vertical line at HR=1 represents no effect.

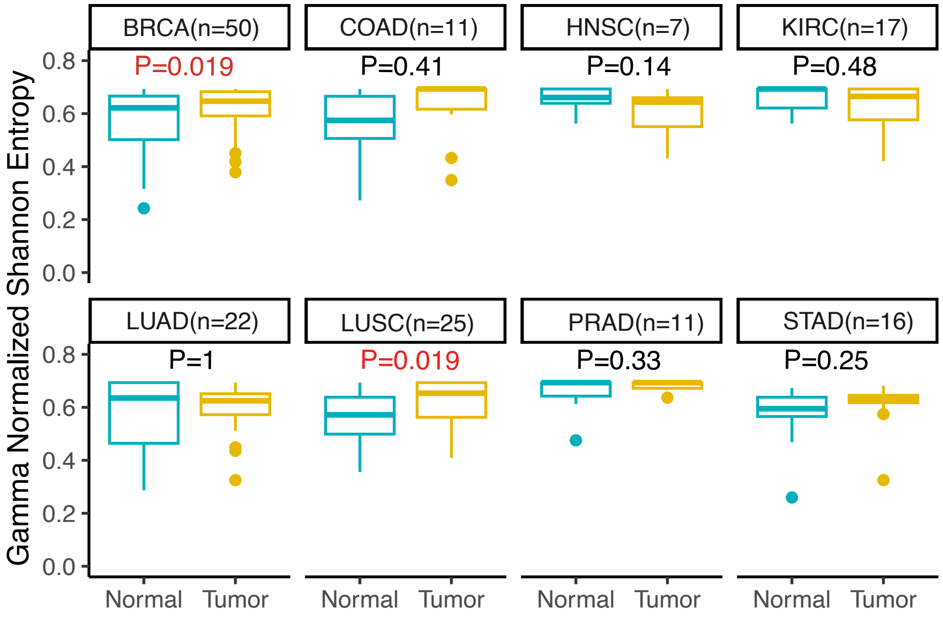


**Supplementary Figure S10.** Diversity metric (Normalized Shannon Entropy) of gamma TCRs between tumor-normal pairs across cancer types. Box represents the median (central line), the 25% and 75% interquartile (IQR) (lower and upper hinges), the ± 1.5 IQR (Tukey whiskers), and outliers (dots). P-value was calculated by two-sided paired Wilcoxon rank-sum test.
